# Characterization of *Streptococcus uberis* Cas9 (SuCas9) – a Type II-A Ortholog Functional in Human Cells

**DOI:** 10.1101/2023.12.11.571105

**Authors:** Polina Selkova, Aleksandra Vasileva, Anatolii Arseniev, Marina Abramova, Olga Musharova, Polina Malysheva, Mikhail Khodorkovskii, Konstantin Severinov

## Abstract

Type II CRISPR-Cas9 RNA-guided nucleases are commonly used for genome engineering. To date, all characterized Cas9-based genome editors, including the widely used SpCas9, have limitations such as their relatively large size and restriction of targets flanked by a specific PAM sequence. Here, we biochemically characterized more compact SpCas9 ortholog, SuCas9, from *Streptococcus uberis*, a bacterium inhabiting the mammary glands of dairy cattle. SuCas9 recognizes a novel 5′-NNAAA-3′ PAM, efficiently cleaves DNA *in vitro*, and is active in human cells. The study of SuCas9 has the potential to expand the range of applications of CRISPR-Cas9 enzymes in medicine and biotechnology.

## 1. Introduction

CRISPR-Cas systems (short for Clustered Regularly Interspaced Short Palindromic Repeats and CRISPR-associated proteins) are bacterial and archaeal immune systems that use RNA-guided Cas nucleases to degrade the genomes of invading entities. These nucleases act as “molecular scissors” that can precisely target and cut specific sections of the intruder’s genetic material, with the RNA component serving as a guide to direct the nuclease to the desired target sequence and ensure efficient degradation of the invading genomes [1].

CRISPR-Cas systems consist of CRISPR arrays and CRISPR-associated *cas* genes. A CRISPR array is a DNA segment in which short repeats (direct repeats; DRs) are separated by unique sequences called spacers. After transcribing the CRISPR array and processing the resulting non-coding RNA, short CRISPR RNAs (crRNAs) are produced. Each crRNA contains a sequence of one of the spacers flanking the repeated fragment. Individual crRNAs bind to Cas nucleases to form effector complexes. When the invading DNA (or RNA, depending on the system) contains a sequence that matches one of the spacers in the CRISPR array, the effector complex identifies it based on complementary interactions with crRNA. The activation of the nuclease activity in the protein component of the effector complex results in the cleavage of the foreign nucleic acid at the recognition site, leading to its eventual degradation [2,3,4].

CRISPR-Cas systems classified as class 2 and type II utilize single-subunit proteins known as Cas9 as effectors [5]. The simplicity of these systems, which require only one protein for programmable nucleic acid cleavage, makes them highly practical and convenient to use. As a result, RNA-guided CRISPR-Cas9 systems have dramatically advanced genome editing. The precise modification of the genetic composition of diverse organisms and cell types through this cutting-edge technology opens up endless possibilities for research and applications in fields such as medicine, agriculture, and biotechnology [6].

Cas9, derived from *Streptococcus pyogenes* (SpCas9), was the first CRISPR-Cas nuclease that enabled targeted mutagenesis in human and animal cells [7,8,9,10]. Despite the availability of several other orthologs, it remains the most widely used Cas9 nuclease for genome editing.

However, all the characterized Cas9-based genome editors, including SpCas9, have limitations. One limitation of SpCas9 is its large size, with 1.368 amino acids (158.46 kDa) [11]. This poses a challenge when attempting to package the *spcas9* gene and its guide RNA sequence together into certain viral vectors, such as adeno-associated virus (AAV), for efficient delivery into cells *in vivo* [12].

To achieve the desired targeting of a specific Cas9 effector, it is necessary to consider not only the complementarity of the RNA guide to its intended sequence but also the presence of a unique Protospacer Adjacent Motif (PAM) sequence in the target genome. PAM flanks the target from the 3′ end and is recognized by the special domain (PAM-interacting domain, PID) of Cas9. PAM limits the number of targets that can be edited [13]. For example, SpCas9 requires 5′-NGG-3′ PAM, Nme1Cas9 – 5′-NNNNGATT-3′ [14], St1Cas9 – 5′-NNAGAAW-3′ [11], Cje1Cas9 – 5′-NNNNRYAC-3′ [11], SaCas9 – 5′-NNGRRT-3′ [12], CdCas9 – 5′-NNRHHHY-3′ [12], GeoCas9 – 5′-NNNNCRAA-3′ [15], CcCas9 – 5′-NNNNGNA-3′ [16], PpCas9 – 5′-NNNNRTT-3′ [17], DtCas9d – 5′-NRC-3′ [18] and CoCas9 – 5′-NRRWC-3′ [19].

To address these challenges, scientists have been investigating novel Cas9 nucleases that are smaller in size and have the ability to recognize alternative PAM sequences. These nucleases with novel properties will enable the expansion of existing genome editing tools and selection of optimal effectors for resolving specific issues.

In this study, we biochemically characterized SuCas9, an effector enzyme from the type II-A CRISPR-Cas system of *Streptococcus uberis* DSM 20569. This bacterium inhabits the mammary glands of dairy cattle and is one of the most frequently identified pathogens causing bovine mastitis [20]. SuCas9 has a smaller size than SpCas9, with 1.126 amino acids and a molecular weight of 131.78 kDa. We demonstrated that SuCas9 is an active nuclease that efficiently cleaves DNA targets *in vitro* across a wide temperature range, from 30°C to 45°C. Also SuCas9 nuclease exhibits its effectiveness in human cells by generating insertions and deletions (indels) in HEK293T genome targets flanked by a novel 5′-NNAAA-3′ PAM. To enhance the potential of SuCas9, we designed a single guide RNA form, created SuCas9 nicking enzymes, and a catalytically “dead” SuCas9 form that can bind DNA but does not cleave it.

## 2. Results

### 2.1 The organization of Streptococcus uberis CRISPR type II-A locus

SuCas9 is an effector endonuclease of the only CRISPR-Cas system that has been bioinformatically identified in the genome of *Streptococcus uberis* DSM 20569. This system was described by Makarova et al. [5] and has been classified as belonging to the II-A type.

The *S. uberis* type II-A CRISPR-Cas locus includes the *cas9* gene, which encodes relatively small effector (1.126 amino acids). Additionally, there are the *cas1* and *cas2* genes, as well as the subtype-specific *csn2* gene, comprising the CRISPR adaptation module. The *S. uberis* CRISPR array contains sixteen 36-bp DRs interspaced with 30-bp spacer sequences. The *cas9* gene is followed by a putative tracrRNA-encoding sequence that is partially complementary to DRs (Figure 1b). *In silico* co-folding of part of DR with the putative tracrRNA predicts stable secondary structures (Figure S6a).

**Figure 1:**
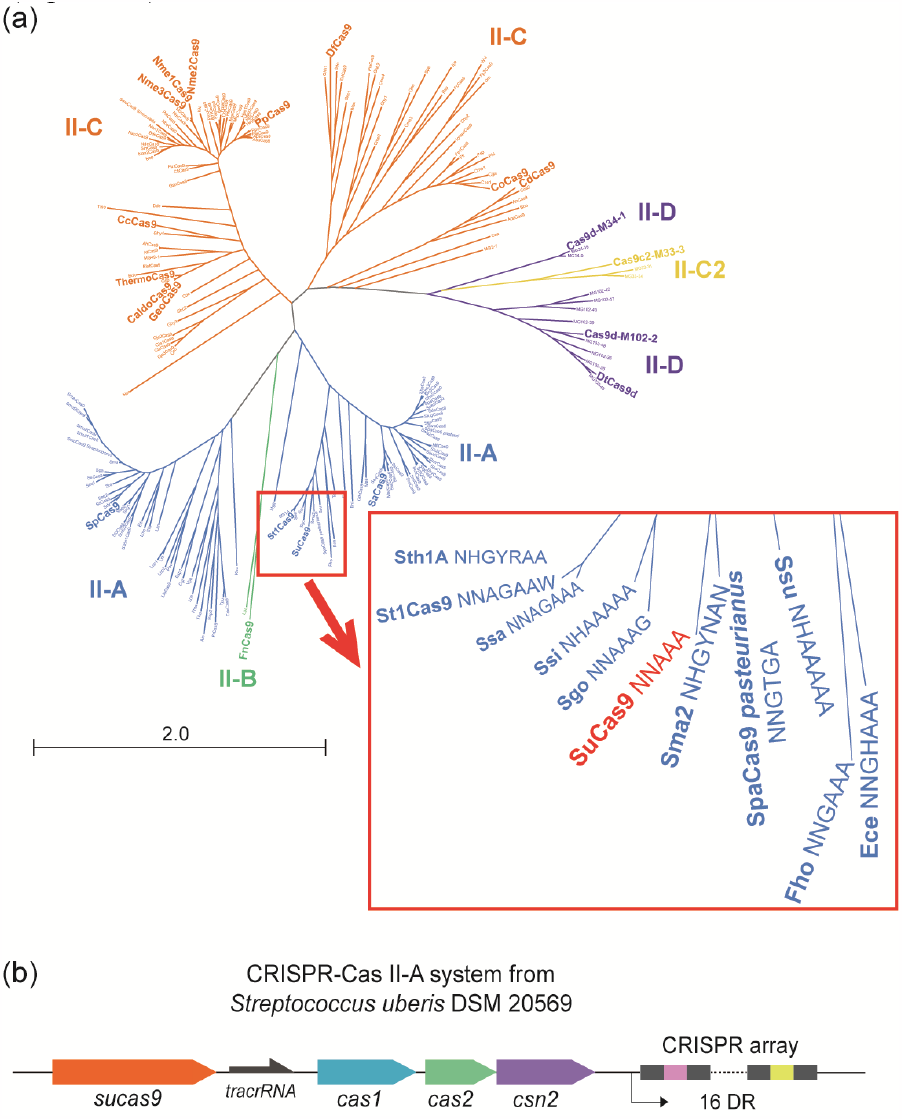
(a) Phylogenetic tree showing the relationship of SuCas9 to other biochemically characterized Cas9 orthologs (see Table S1). Blue, green, orange, yellow and purple tree branches indicate whether the effector belongs to type II-A, type II-B, type II-C, type II-C2 or type II-D. (b) Organization of the *Streptococcus uberis* DSM 20569 type II-A CRISPR-Cas system (positions 1 118 642 – 1 1117 34 of the *S. uberis* genome sequence (NZ_LS483397.1)).

### 2.2 SuCas9 PAM requirements

To investigate SuCas9 PAM sequence, we carried out *in vitro* PAM screening experiments. The purified recombinant SuCas9 was derived from *E. coli* Rosetta (DE3) cells (Figure S1), and the crRNAs and tracrRNAs were synthesized *in vitro* based on bioinformatics predictions. To evaluate the nuclease activity of SuCas9, we tested its ability to cleave linear DNA PAM libraries that contained 20-bp target site flanked with seven randomized nucleotides at the 3′ end. The sequences of the libraries used are listed in Table S4. Uncleaved molecules were purified after agarose gel electrophoresis and sequenced using an Illumina platform. Reactions without tracrRNAs were used as the negative controls.

Initially, 400 nM of SuCas9, along with crRNA and tracrRNA, was exposed to the PAM library at 37°C for 30 min. A total of 16 384 unique PAM sequences were found for the depleted and control samples. The median coverage of every individual PAM variant comprised 388 and 534 reads for the depleted and control samples, respectively. The comparison of PAM sequences representation in the SuCas9 depleted sample and the negative control allows one to determine the PAM preferred by SuCas9. A total of 659 PAM sequences were statistically significantly depleted. The PAM logo representation showed the importance of “A” in the 3^rd^, 4^th^, and 5^th^ positions flanking the protospacer (Figure 2a). The wheel approach for PAM representation developed by Leenay et al. [21] revealed the additional preference “G” in the 5^th^ PAM position (Figure 2b).

**Figure 2:**
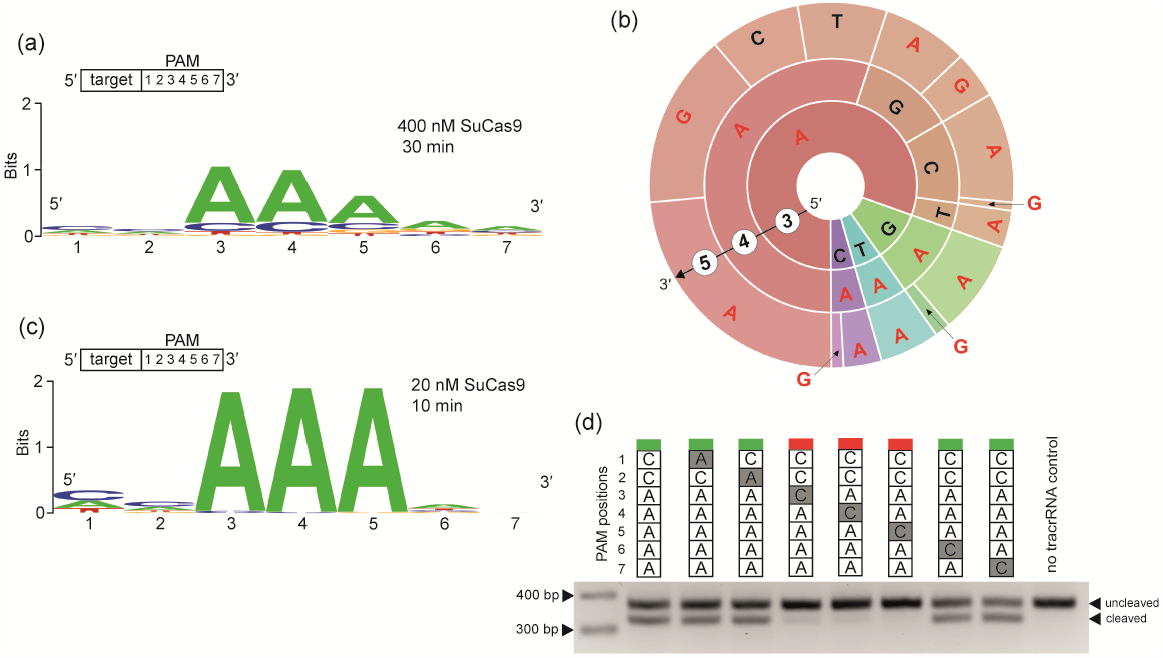
(a) SuCas9 PAM logo determined by an *in vitro* PAM screening experiment. Samples were incubated with 400 nM SuCas9 at 37°C for 30 min. (b) PAM wheel representation of the panel “a” *in vitro* PAM screening experiment results for the 3^rd^, 4^th^, and 5^th^ nucleotide positions of PAM. Nucleotide positions from the inner to the outer circle match the PAM positions moving away from the protospacer. The proportion of the sector’s area in the PAM wheel corresponds to the relative depletion in the library and, as a result, to the significance of the nucleotide. Colors are used to keep track of the PAM sequence, depending on the nucleotide in the first position. (c) SuCas9 PAM logo determined using an *in vitro* PAM screening experiment. Samples were incubated with 20 nM SuCas9 at 37°C for 10 min. (d) Single-nucleotide substitutions in the 3^rd^, 4^th^, and 5^th^ positions of PAM prevent SuCas9 from cleaving DNA. Samples were incubated with 20 nM SuCas9 at 37°C for 10 min. An agarose gel displaying the outcomes of electrophoretic separation of cleavage products of targets with PAM sequences presented at the top. The color labels at the top of PAM sequence columns (green or red) indicate whether cleavage took place.

These results reflect a preference of SuCas9 for PAM sequences only at high protein concentrations *in vitro*. To improve the accuracy of our SuCas9 *in vitro* PAM predictions for use *in vivo*, we conducted additional *in vitro* PAM screening experiments using smaller concentrations of nuclease and a shorter incubation time of 10 minutes. As a result, we found that the conservation of “A” at the 3^rd^, 4^th^ and 5^th^ positions of PAM increased (Figure 2c, Figure S2).

The results of additional PAM screening experiments with DNA library of different protospacer sequence confirmed the previous results and emphasized the importance of the 3^rd^, 4^th^ and 5^th^ PAM positions (Figure S3).

Based on the results presented above, we concluded that the most preferred SuCas9 PAM sequence corresponds to the following consensus: 5′-NNAAANN-3′.

To validate the preferences of SuCas9 for PAM sequences, we made single-base changes to the inferred consensus PAM sequence located adjacent to a 5′-TATCTCCTTTCATTGAGCAC-3′ target site on the 3′-end. The use of primers with modified PAM-encoding sequences allowed for the PCR-based synthesis of 374-basepair DNA cleavage substrates. The cleavage of these fragments by SuCas9 should result in the generation of a shorter fragment, reduced by approximately 49 base pairs, which can be easily distinguished from uncleaved DNA through agarose gel electrophoresis. Initially, we incubated DNA substrates with different PAM sequences with 400 nM of SuCas9 for 30 min. This experiment confirmed our earlier assumption that this nuclease is highly active *in vitro* and, therefore, has a reduced requirement for PAM sequences under such conditions (Figure S4). To improve the accuracy of PAM sequence detection, we reduced the concentration of SuCas9 to 20 nM and shortened the incubation time to 10 minutes. Under these conditions, substitutions at 1^st^, 2^nd^, 6^th^ and 7^th^ positions did not affect cleavage (Figure 2d, Figure S4), while the substitution of “A” in the 3^rd^, 4^th^ and 5^th^ positions abolished SuCas9 nuclease activity.

Next, we tested whether SuCas9 could be targeted to different sequences flanked by 5′-NNAAA-3′ (Figure 3). Sixteen unique 20 bp sequences, each flanked by different PAM variants, were selected from a 1456 bp linear DNA fragment (PCR product derived from the *GRIN2b* human gene). The Table S6 provides information about the sequence and position of SuCas9 target sites on this fragment. 20 nM of SuCas9 was incubated with this dsDNA fragment, tracrRNA, and crRNAs complementary to one of the 16 targets for 10 min at 37°C. Almost all nuclease targets that were effectively cleaved had a flanking sequence that matched the PAM consensus 5′-NNAAA-3′. However, some of the targets whose PAM sequences differed from the consensus by one letter (targets 1, 10, 11, and 13) were also cleaved. Nevertheless, the last two targets were cleaved with a very low efficiency. The probable reason for the recognition of these targets by SuCas9 nuclease is its high *in vitro* activity, which may not accurately reflect the *in vivo* situation. Overall, SuCas9 failed to cleave nine out of the twelve targets that were flanked by sequences that differed from the PAM consensus (Figure 3). These results demonstrate that SuCas9 can recognize and cleave specific DNA targets that are flanked by 5′-NNAAA-3′ PAMs *in vitro*.

**Figure 3.**
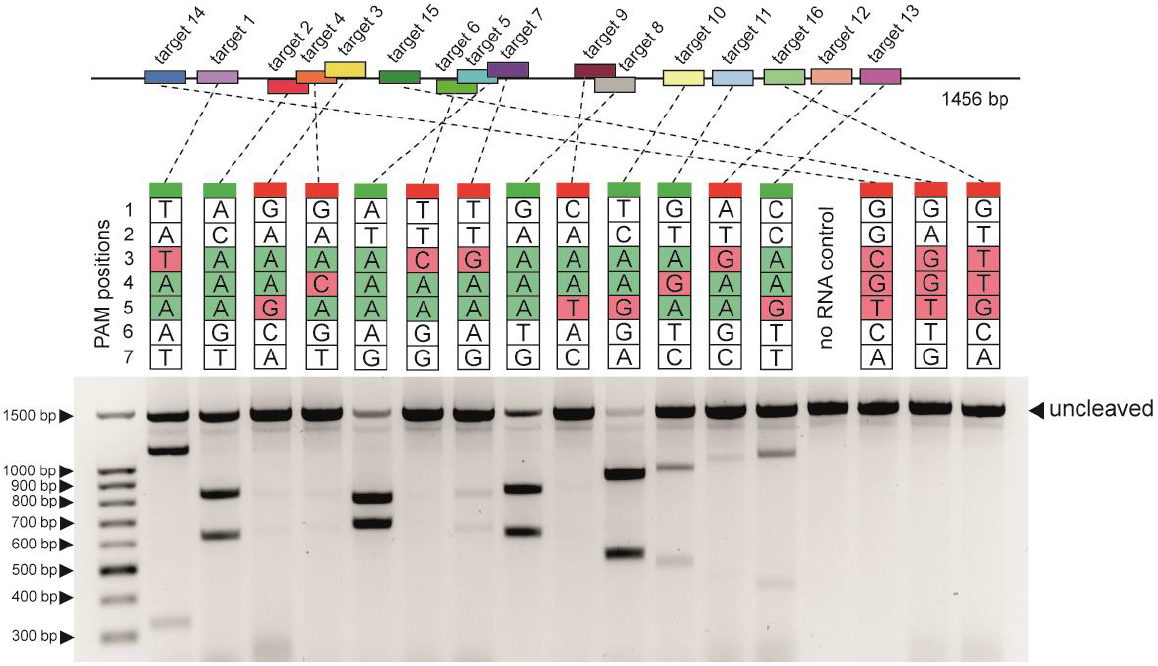
SuCas9 cleaves different 20 bp DNA targets flanked by a 5′-NNAAA-3′ PAM. Sixteen different 20 bp sequences flanked by unique PAMs in a 1456 bp linear DNA fragment were interrogated by SuCas9, tracrRNA, and individual crRNAs complementary to each target. The scheme above shows the positions of different target sites in the 1456 bp fragment. A gel showing results of *in vitro* cleavage of targets with indicated PAMs is presented below. The color labels at the top of PAM sequence columns (green or red) indicate whether cleavage took place. Expected sizes of DNA cleavage products can be found in Table S6.

To determine SuCas9 PAM requirements we also used the CRISPRTarget tool [22] to predict the putative targets (protospacers) of *S. uberis* CRISPR array spacers. We selected only sequences from *Streptococcus* phages for analysis. Although the results were not sufficient to generate logo for PAM, which could be considered reliable (Table S7), they correlated with the *in vitro* PAM logo 5′-NNAAA-3′ and may further confirm its validity.

Overall, we conclude that SuCas9 nuclease specifically cleaves DNA targets flanked with 5′-NNAAA-3′ PAM sequence at the 3′ side of protospacers.

### 2.3 SuCas9 temperature preferences

The range of optimal working temperatures is an important factor for determining the application of Cas nucleases. It is crucial to consider this when using Cas nucleases in various applications because the working temperature can greatly affect their efficiency. The temperature dependence of SuCas9 nuclease activity was determined by testing its performance on the target flanked by 5′-NNAAA-3′ PAM on either a linear DNA fragment or a plasmid. SuCas9 efficiently cleaved the plasmid DNA within a temperature range of 30–45°C and demonstrated the same efficiencies for cleaving linear and supercoiled DNA (Figure S5). Based on this, it can be inferred that SuCas9 has a relatively broad working temperature range, making it a versatile option for multiple applications.

### 2.4 SuCas9 sgRNA design

To improve the utility of SuCas9 in biotechnology, we created a single-guide RNA (sgRNA) that linked the DR segment of crRNA to tracrRNA, similar to SpCas9 and other Cas9 orthologs [9]. We tested two sgRNAs with 36- and 31-nt DR-derived segments *in vitro* (Figure S6b) and found that SuCas9 cleaved target dsDNA more efficiently when in complex with sgRNAs than when it was combined with individual crRNA and tracrRNA (Figure S6c). The shorter sgRNA2 was chosen for further experiments.

### 2.5 SuCas9 mediates genome editing in human cells

Next, we investigated whether SuCas9 was capable of genome editing in human cells. Codon-optimized SuCas9, PpCas9, and SpCas9 (the latter two are positive controls) were cloned into plasmid vectors under the regulation of the constitutive CMV promoter. A GFP coding sequence was fused in-frame with nuclease open reading frames through a sequence encoding a P2A self-cleaving peptide. Appropriate sgRNA coding sequences were introduced into the plasmids upstream of the nuclease genes under the control of the U6 promoter. Cas9 nucleases were targeted to the human *EMX1* gene (Table S8). Changing the spacer length of the guide RNA is known to have a significant effect on interference efficiency [12,17,23]. Given this, for each SuCas9 site we used sgRNAs with spacer sequences of two lengths, 20 and 22 nt. For PpCas9 and SpCas9, only one spacer length with a proven efficiency was used.

The resulting plasmids were transfected into the HEK293T cells (Figure 4a). The production of the recombinant SuCas9 protein was confirmed by western blot analysis (Figure S7).

**Figure 4:**
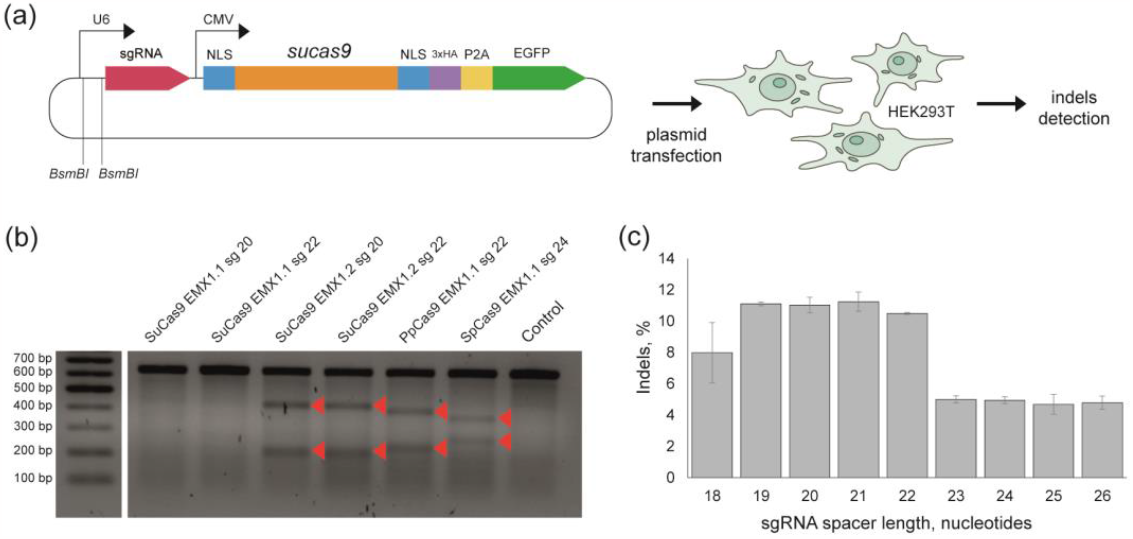
(a) Schematic of SuCas9-mediated genome disruption in HEK293T cells. Above: scheme showing the design of a plasmid used for *sucas9* gene and sgRNA expression. The *sucas9* gene indicated by an orange rectangle. The CMV and U6 promoters are indicated by black arrows. The plasmid was transfected into HEK293T cells, and genomic DNA was extracted for indel frequency assessment through an *in vitro* assay with T7 endonuclease I or high-throughput sequencing of the targeted region.(b) T7 endonuclease I indel detection assay showing SuCas9-mediated cleavage of the *EMX1* gene in the HEK293T genome. Products of target DNA fragment incubation with T7 endonuclease I are indicated by red triangles. (c) Influence of sgRNA spacer length on SuCas9-mediated indel formation efficiency in EMX1.2 target site. HEK293T cells were transfected with SuCas9 sgRNA plasmids encoding sgRNAs with spacer segments of various lengths. Mean values and standard deviations obtained from three biological replicates are shown.

Three days after transfection, genomic DNA was extracted from a heterogeneous population of modified and unmodified cells and indel formation was assessed using the T7 endonuclease I detection assay. SuCas9 introduced indels in the EMX1.2 target, but failed to modify the EMX1.1 target. Where cleavage was observed, sgRNAs with spacer sequences of 20 and 22 nt showed similar editing efficiencies (Figure 4b).

The same targets on *EMX1* and an additional target on *GRIN2b* gene (Table S8) were used for further indel frequency analysis by high-throughput sequencing. Although the T7 endonuclease I detection assay failed to detect modifications of EMX1.1, indels were detected for all three targets based on NGS results (Figure S8).

Target EMX1.2 was selected for further evaluation of SuCas9 DNA modification efficiency.

The length requirements of the SuCas9 sgRNA spacer for optimal genome editing were further investigated. HEK293T cells were transfected with SuCas9 carrying plasmids similar to those described above, but coding for sgRNAs of different spacer lengths targeting the EMX1.2 site in the human genome. The efficiency of DNA modification was assessed using high-throughput sequencing of the target site. The results showed that SuCas9 efficiently introduced double-stranded breaks in genomic DNA with sgRNAs of 19-21 nt spacer length (Figure 4c). Equally efficient cleavage with three different spacer lengths may open up a wider range of possibilities for selecting guide RNAs for specific genomic targets.

### 2.6 The specificity of SuCas9

The use of Cas nucleases in biotechnology requires the recognition of specific targets. To investigate the cleavage specificity of SuCas9 in eukaryotic cells, we performed a preliminary study. Our on-target was the DNA site in *EMX1*, which SuCas9 had efficiently cleaved in previous experiments (EMX1.2). HEK293T cells were transfected with plasmids carrying the SuCas9 genome editing system targeting this site. In this experiment, we directed SuCas9 to on-target sequences using sgRNAs with a spacer length of 20 nt. Such sgRNAs provide sufficient DNA cleavage efficiency and allow the identification of more potential off-target sequences than sgRNAs with longer spacer segments. Three days post-transfection genomic DNA was extracted and indel frequency at the on-target as well as at likely off-target sites (sequences differing by up to 3 nt from on-target site) was assessed by targeted amplicon high-throughput sequencing. Computational analysis using CRISPResso2 [24] indicated significant modifications compared to the negative controls only at on-target site (Figure 5). The modification of the remaining off-target sites compared to the negative control was statistically insignificant. Based on these preliminary results, we conclude that SuCas9 can be considered a promising nuclease in terms of its specificity, although additional studies using Digenome-seq [25], GUIDE-seq [26], BLESS [27], BLISS [28], or other methods should be used to fully estimate its off-targeting activity.

**Figure 5.**
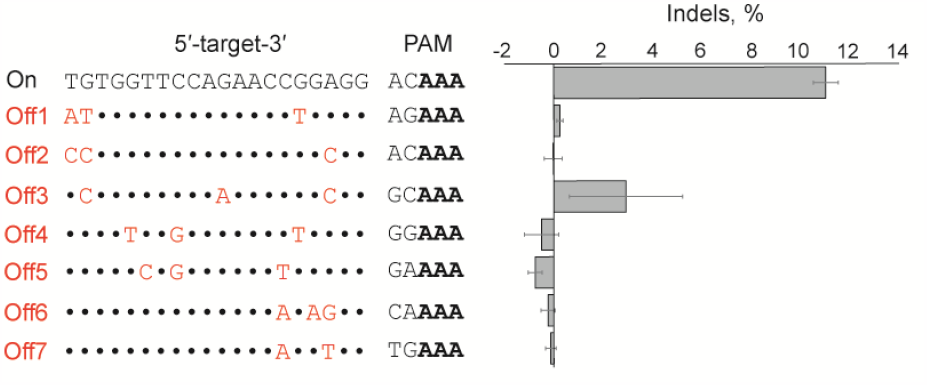
Specificity of genomic DNA cleavage using SuCas9. The indel frequency at the EMX1.2 target as well as at the corresponding off-target sequences was assessed by targeted amplicon sequencing of genomic DNA of HEK293T cells transfected with plasmids carrying the SuCas9 genome editing system. Left: sequences of on- and off-target sites are shown. Mismatches in the off-target sequences are shown in red. Right: Frequency of indel formation at each site, mean values and standard deviations obtained from three biological replicates are shown.

### 2.7 SuCas9 nickases creation

We have also created SuCas9 versions that nick DNA, as well as “dead” SuCas9 that binds DNA but does not cleave it [9,29]. By comparing the amino acid sequences of SuCas9 with those of SpCas9, we were able to identify the locations of the active sites in the RuvC and HNH domains (Figure S9). We then individually mutated these sites, resulting in the inactivation of the corresponding nuclease domains. The D10A mutation inactivated the RuvC domain and the H600A mutation inactivated the HNH domain, causing the SuCas9 mutants to behave as nickases. Finally, we created a double mutant that was catalytically inactive (Figure S1, Figure S10).

### 2.8 Phylogenetic relationship of SuCas9

On the phylogenetic tree of Cas9 orthologs SuCas9 is located on a separate branch of type II-A enzymes (Figure 1a). The Cas9 enzymes with the highest similarity to SuCas9 were Sma2 from *Streptococcus macedonicus* 33MO (130.87 kDa) and SpaCas9 from *Streptococcus pasteurianus* ATCC 43144 (131.08 kDa). Both orthologs differ significantly from SuCas9 in terms of the recognizable PAM sequence: Sma2 – 5′-NHGYNAN-3′ [30], SpaCas9 – 5′-NNGTGA-3′ [12]. Additionally, slightly more distinct SuCas9 orthologs that recognize similar PAM sequences were discovered in the same phylogenetic branch: Ssi – 5′-NHAAAAA-3′, Sgo – 5′-NNAAAG-3′, Ssu – 5′-NHAAAAA-3′. These nucleases have also been detected in various species of streptococci. However, nuclease activity has only been demonstrated *in vitro* for Ssi, Sgo, and Ssu, there is currently no information available on their activity in human or animal cells [30].

## 3. Discussion

Despite the extensive use of Cas9 nucleases for genome engineering, to date, only a few Cas9 orthologs have been considered well characterized. Given the diversity of type II CRISPR–Cas systems, Cas9 orthologs can show significant variations in PAM requirements, specificity, and other biochemical properties. Moreover, specificity and small size are crucial factors in the use of genome engineering tools, as their genes can be delivered in eukaryotic organisms using size-restricted vectors, such as AAV.

Recently, there have been notable attempts to modify PAM recognition, loosen PAM requirements, shorten nucleases, and develop hybrid forms for commonly spread Cas nucleases using protein engineering. However, concerns have been raised regarding the specificity and efficiency of target recognition by PAM-free Cas nucleases. Another approach is to create a collection of Cas nucleases with different PAM requirements, which can be used for any target. To build a complete collection, it is essential to systematically search for natural Cas9, Cas12, and other Cas orthologs and characterize them. Furthermore, when selecting nucleases for gene therapy, it is particularly crucial to identify those that are highly expressed in eukaryotic cells. This capability is essential for effective application in gene engineering, but not all nucleases possess this capability. Therefore, biochemical characterization of different Cas enzymes is essential for creating a comprehensive PAM-flexible Cas nuclease toolbox with minimal PAM limitations, small size, and expression in eukaryotic cells.

In this study, we performed biochemical characterization of SuCas9 from the *Streptococcus uberis* type II-A CRISPR-Cas system, both *in vitro* and *in vivo*. Our findings demonstrated that SuCas9 displays remarkable efficiency and specificity, especially when targeting simple 3-nucleotide PAM sequences (5′-NNAAA-3′) in eukaryotic cells. This makes SuCas9 an ideal and dependable platform for a wide range of biotechnological applications. In addition, we developed catalytically inactive SuCas9, which can be used as a tool for targeted gene activation/inactivation, live-cell imaging, and base editing.

Although the specificity of SuCas9 should be thoroughly investigated and analyzed in future research, the data acquired in this study strongly indicated that the nuclease exhibits minimal off-targeting activity, making it a potentially promising tool for genetic engineering applications.

## 4. Materials and Methods

### 4.1 Plasmid construction

For pET21a_SuCas9 plasmid cloning, SuCas9 coding sequence was PCR amplified with locus_ SuCas9_F and locus_SuCas9_R primers using *Streptococcus uberis* genomic DNA (DSM 20569, https://www.dsmz.de/collection/catalogue/details/culture/DSM-20569) as a template. The resulting fragment was then inserted into *XhoI* and *NdeI* digested pET21a vector by NEBuilder HiFi DNA Assembly Cloning Kit (NEB, E5520). As a result, SuCas9 contained His6 on C-termini. The link to the vector map and primers are presented in Tables S2 and S3, respectively.

For expression in human cells SuCas9 gene was codon optimized and inserted into plasmid under regulation of CMV promoter. sgRNA expression was driven by U6 promoter. The vector map is presented in the Table S2.

To generate the plasmids encoding SuCas9_D10A, SuCas9_H600A nickase forms, and the “dead” form dSuCas9(D10A, H600A), the pET21a_SuCas9 vector was modified through PCR amplification of the entire vector using PCR primers that included the desired substitutions in their sequences. The circularization of the plasmids was then achieved using the NEBuilder HiFi DNA Assembly Cloning Kit (NEB, E5520). The primers utilized for site-specific mutagenesis are presented in Table S2, and the links to the mutated plasmid vector maps are provided in Table S2.

### 4.2 Protein expression and purification

For recombinant SuCas9 purification, competent *E. coli* Rosetta (DE3) cells were transformed with pET21a_SuCas9 plasmid and grown until OD_600_ = 0.4-0.6 in 500 ml LB media supplemented with 100 μg/ml ampicillin. The target protein synthesis was induced by the addition of 1 mM isopropyl-β-D-1-thiogalactopyranoside (IPTG). After 12 h of growth at 18°C, cells were centrifuged at 3 500 g for 30 min. The pellet was resuspended in lysis buffer (50 mM Tris (pH 7.5 at room temperature), 500 mM NaCl, 10 mM imidazole, and 1 mM β-mercaptoethanol) supplemented with 1 mg/ml lysozyme (PanReac AppliChem, A3711,0010), and cells were lysed by sonication. The cell lysate was centrifuged at 16 000g (4°C) for 1 h and filtered through a syringe filter with a cellulose nitrate membrane (pore size 0.4 µm). The lysate was applied to a 1 ml HisTrap HP column (GE Healthcare) and eluted with 300 mM imidazole in the same buffer without lysozyme. After affinity chromatography, fractions containing the nuclease were applied to a Superdex200 Increase 10/300 GL (GE Healthcare) column equilibrated with a buffer containing 50 mM Tris–HCl (pH 7.5 at room temperature), 500 mM NaCl, and 1 mM 1,4-Dithiothreitol (DTT). Fractions containing SuCas9 monomer were pooled and concentrated using a 30 kDa Amicon Ultra-4 centrifugal unit (Merc Millipore, UFC803008). Glycerol was added to a final concentration of 10%, and samples were flash-frozen in liquid nitrogen and stored at -80°C. Purity of the nucleases was assessed by denaturing 10% polyacrylamide gel electrophoresis.

To obtain recombinant SuCas9_D10A, SuCas9_H600A nickase forms, and the “dead” form dSuCas9, the same protocol was used.

### 4.3 In vitro DNA cleavage assay

DNA cleavage reactions were performed using the recombinant SuCas9 protein and linear dsDNA targets, which were PCR products of human *GRIN2b* gene amplification. The reaction conditions were as follows: 1× CutSmart buffer (50 mM Potassium Acetate, 20 mM Tris-acetate, 10 mM Magnesium Acetate, 100 µg/ml BSA or recombinant albumin; NEB), 0.5 mM 1,4-Dithiothreitol (DTT), 40 nM DNA, recombinant protein (concentration varied depending on the experiment and ranged from 20 nM to 400 nM), and crRNA + tracrRNA or sgRNA (RNA concentrations were 10 times higher than protein concentration). The samples were incubated at 37°C; incubation times are specified separately for each individual experiment. To stop the reaction, 4× loading dye containing 10 mM Tris–HCl (pH 7.8), 40% glycerol, 40 mM ethylenediaminetetraacetic acid, 0.01% bromophenol blue, and 0.01% xylene cyanol was added. The reaction products were then analyzed by electrophoresis in 2% agarose gels. Ethidium bromide pre-staining was used for band visualization on agarose gels.

The *in vitro* SuCas9 PAM screenings were carried out using linear DNA 7N PAM libraries that were created through PCR amplification of a 374 bp or a 359 bp region of the human *GRIN2b* gene. The forward primer used in the amplification process contained seven randomized nucleotides upstream of its 3′-end segment, which was complementary to the target sequence (Table S3, Table S4). 100 nM linear DNA 7N PAM library was incubated with 400 nM, 200 nM, 100 nM or 20 nM recombinant protein, crRNA + tracrRNA (RNA concentrations were 10 times higher than protein concentration). Reactions without tracrRNA were used as negative controls. The reaction was performed at 37°C; incubation times are specified separately for each individual experiment. Reaction products were separated by electrophoresis in agarose gels. Uncleaved DNA fragments were extracted from the gel using the Cleanup Standard kit (Evrogen, BC04). These fragments were used as the templates for two-step PCR. For the first PCR step primers combining target-specific sequences and Illumina adaptor overhangs were used. The result amplicons were used as templates in the second PCR step. During this step 8N barcodes and flow cell linker adaptors were introduced into the sequences (Table S3). The products of the second round PCR were loaded on agarose gel electrophoresis. The PCR fragments were gel-extracted using the Cleanup Standard kit (Evrogen, BC04) and sequenced using Illumina NextSeq, MiSeq, or NovaSeq 6000 platform. The sequencing was performed with either single-end or paired-end reads, and the number of cycles used was either 150 or 300.

For testing SuCas9 activity at different temperatures a mix of the SuCas9 protein with *in vitro* synthesized crRNA and tracrRNA in the buffer, and the target DNA, also in the cleavage buffer, were first incubated separately at the chosen temperature for 2 min, and then were combined and incubated for additional 10 min at the same temperature. The following concentrations were used: 12 nM DNA, 240 nM SuCas9, 1.2 µM crRNA, 1.2 µM tracrRNA – for *in vitro* cleavage of a linear DNA fragment; 4 nM DNA, 80 nM SuCas9, 400 nM crRNA, 400 nM tracrRNA – for *in vitro* cleavage of a plasmid DNA.

To test the nuclease activity of the SuCas9 mutants, DNA cleavage reactions were performed under the same conditions as above using 20 nM target DNA, 200 nM wild-type or mutant SuCas9, and 1 μM sgRNA. The samples were then incubated at 37°C for 30 min. Proteinase K (Thermo Scientific, EO0491) was added to inactivate the effector (1 μl per sample, incubation for 15 min at 37°C). Next, the reactions were divided into two halves and the DNA cleavage products were analyzed by gel electrophoresis under native (1.5% agarose gel electrophoresis) and denaturing conditions (denaturing 5% polyacrylamide gel electrophoresis, 8 M urea).

### 4.4 Cell culture and plasmid transfection

HEK293RT cells were maintained in Dulbecco’s modified Eagle’s medium (DMEM) supplemented with 10% fetal bovine serum at 37°C with 5% CO_2_. Cells were seeded into 24-well plates (Eppendorf) one day prior to transfection. Cells were transfected using Lipofectamine 2000 (Thermo Fisher Scientific, 11668019), following the manufacturer’s protocol. For each well of a 24-well plate a total of 500 ng plasmids was used. Three days after transfection, the cells were harvested, and genomic DNA was extracted using QuickExtract solution (Lucigen, QE0950).

### 4.5 Western blotting analysis

72 hours post-plasmid transfection, HEK293T cells cultured in 24-well plates were lysed in protein lysis buffer on ice for 15 min and then gently sonicated. Samples were boiled for 5 min and loaded onto a denaturing 10% polyacrylamide gel according to Laemmli [31] and run in 1X TBE buffer. The gel was transferred to a nitrocellulose membrane in 1X Transfer Buffer at 100V for 1 h on ice. The blot was incubated in primary antibody overnight (1:1000 anti HA-tag HA.C5). Then it was washed three times in PBS-T to remove excess primary antibody and incubated in secondary antibody (1:20.000 Anti-Mouse IgG) for one hour at room temperature. The blot was then washed three times in PBS-T, incubated in SuperSignal WestPico Chemiluminescent Substrate (Thermo), and developed on the ChemiDoc digital imager (BioRad).

### 4.6 Assessment of indel efficiency

Genomic DNA was obtained from the transfected cells as described in “2.4. Cell culture and plasmid transfection”.

For the T7 endonuclease I indel detection assay, the genomic region surrounding the CRISPR target site was PCR amplified using the primers listed in Table S3. The PCR fragments were gel-extracted using a Cleanup S-Cap kit (Evrogen, BC04). Next, an indel detection assay was performed using T7 endonuclease I (Genes2Me, TEND-250U) according to the manufacturer’s instructions.

For indel frequency analysis by high-throughput sequencing, the genomic region surrounding the CRISPR target site was amplified using a two-step PCR. First, primers that combined target-specific sequences and Illumina adaptor overhangs were used. The second PCR was performed using first-round amplicons and the same primers as those used for the *in vitro* PAM screening (Table S3).

Second-round PCR products were subjected to agarose gel electrophoresis. PCR fragments were gel-extracted using the Cleanup Standard kit (Evrogen, BC04) and sequenced using were sequenced using NovaSeq 6000 with 300 paired-end cycles.

### 4.7 Computational sequence analysis

For the analysis of SuCas9 *in vitro* PAM screening results, Illumina reads were filtered by requiring an average Phred quality (Q score) of at least 20. The resulting reads were mapped against the corresponding reference sequence using BWA [32]. All unmapped reads were discarded from the analysis. The degenerate seven-nucleotide region was extracted from the sequences. WebLogo [33] was used to generate a logo based on statistically significant (one-sided Pearson chi-square test with a P-value less than 10^™7^) depleted PAM sequences. The PAM wheel was constructed according to Leenay et al. [21]. The PAM wheel is a convenient visualization of a small number of PAM positions (up to four). The PAM wheel was constructed for SuCas9 PAM positions that were found to be significant according to the PAM logo. For this purpose, 3^rd^, 4^th^ and 5^th^ PAM region positions were extracted from sequences resulting in a 3-nucleotide long set of PAM variants. The occurrence frequency of each PAM variant was recalculated and used for PAM wheel construction. The depletion values of three nucleotide sequences (3^rd^, 4^th^ and 5^th^ PAM positions) for PAM wheel were counted similarly to Maxwell et al. [34]: D_coef_ = log_2_[[total N sample/total N control] ^*^[N PAM control/N PAM sample]], where N is a number of reads. PAM sequences with a positive depletion coefficient were used as input for PAM wheel construction.

SuCas9 spacers were analysed using the CRISPRTarget tool [22] to identify potential bacteriophage protospacer sequences. A maximum of four mismatches between the spacers and putative protospacer sequences were considered a strong protospacer match. The seven nucleotide regions downstream of unique putative protospacers were used for further analysis.

Indel frequencies were estimated using CRISPResso2 analysis package [24]. A window of 20 bp around the target site and quantification window center corresponding to 3 nt from the 3′ end of the guide were provided to detect possible mutations. Fragments derived from the untransfected cells were used as controls. Analysis of control target fragments allows for the determination of the approximate percentage of modifications introduced due to errors accumulated during PCR and subsequent sequencing. The resulting indel percentage was calculated as [indel % in transfected cells] – [indel % in untransfected cells] for a certain region of genomic DNA.

COSMID tool [35] was used to predict the off-target sites. The number of permitted substitutions within the potential target did not exceed 3, and insertions and deletions were not taken into account in the analysis. Off-target indel efficiency was measured through next-generation sequencing analysis. The resulting sequencing data were analysed employing the CRISPResso2 tool as described previously for on-target sites.

Phylogenetic tree showing the relationship of SuCas9 to previously described Cas9 orthologs was constructed using Maximum likelihood (ML) method in MEGA 11 [36].

## 5. Patents

RU2804422C1 “Genomic DNA editing system of an eukaryotic cell based on the nucleotide sequence encoding the SuCas9NLS protein”.

